# HES1 protein oscillations are necessary for neural stem cells to exit from quiescence

**DOI:** 10.1101/2021.02.17.431655

**Authors:** Elli Marinopoulou, Nitin Sabherwal, Veronica Biga, Jayni Desai, Antony D. Adamson, Nancy Papalopulu

## Abstract

Quiescence is a dynamic process of reversible cell-cycle arrest. High-level sustained expression of the HES1 transcriptional repressor, which oscillates with an ultradian periodicity in proliferative neural stem cells (NSCs), is thought to mediate quiescence. However, it is not known whether this is due to a change in levels or in dynamics. Here, we induce quiescence in NSCs with BMP4, which does not increase HES1 level, and we find that HES1 continues to oscillate. To assess the role of HES1 dynamics, we express sustained HES1 under a moderate-strength promoter, which overrides the endogenous oscillations while maintaining the total HES1 level within physiological range. We find that sustained HES1 does not affect proliferation or entry into quiescence, however, exit from quiescence is impeded. Thus, oscillatory expression of HES1 is specifically required for NSCs to exit quiescence, a finding of potential importance for controlling reactivation of stem cells in tissue regeneration and cancer.

## Introduction

Quiescence is defined as a state of reversible growth arrest in which cells are not actively dividing but they retain the capacity to re-enter the cell-cycle when exposed to the appropriate signals. It is a common feature adapted by many stem cell populations that enables them to retain a “reservoir” of cells that can replenish actively dividing stem cells that have been depleted and therefore prevent exhaustion of the stem cell pool (van Velthoven and Rando, 2019). As such, dysregulation of this state has huge implications in the health of an organism as in tissue homeostasis and regeneration (Marescal and Cheeseman, 2020). In the adult vertebrate brain neural stem cells (NSCs) exist primarily in two main brain regions, the subgranular zone (SGZ) of the dentate gyrus in the hippocampus and the ventricular-subventricular zone (V-SVZ) which lines the lateral ventricles, where they remain largely in a quiescent state (Doetsch et al., 1999; Doetsch, 2003; Seri et al., 2001; Urban et al., 2019; Obernier and Alvarez-Buylla, 2019). Although adult NSCs continuously generate new neurons *in vivo*, this does not seem to be a sufficient mechanism for regeneration and brain repair. It appears that the extracellular environment can restrict the activation of quiescent NSCs or their full differentiation potential upon activation (Magnusson and Frisen, 2016). Therefore, understanding how quiescent cells can be reactivated is the first key step to regeneration.

The Notch signaling pathway has been shown to play an important role both during embryonic neural development and adult neurogenesis. Deletion of the RBPJ transcription factor, the major effector of Notch signaling, is embryonic lethal at E9.5 and embryos exhibit delayed neural development (de la Pompa et al., 1997). On the other hand, conditional deletion of RBPJ in the adult brain leads to a transient burst in proliferation of NSCs which soon get depleted followed by an eventual loss of neurogenesis (Imayoshi et al., 2010; Ehm et al., 2010). The Notch effector and transcriptional repressor *Hes1* and the proneural gene *Ascl1*, which is regulated by HES1, they also play important roles during neural development. HES1 represses its own transcription which causes HES1 to oscillate with a 2-3h periodicity in NSCs. In turn, ASCL1 also oscillates in anti-phase with a similar period (Imayoshi et al., 2013). Upon differentiation towards neurons, HES1 expression is lost and ASCL1 oscillations switch to sustained expression which induces neuronal differentiation (Imayoshi et al., 2013). Interestingly, light-induced oscillatory *Ascl1* expression in *Ascl1-/-* embryonic NSCs promotes NSC proliferation whereas sustained expression at similar levels enhances neuronal differentiation, highlighting that the pattern of expression only, but not levels, can dictate cell fate (Imayoshi et al., 2013). Similarly, in the adult brain, the ASCL1 dynamic expression controls the transition from NSC quiescence to activation and differentiation towards neurons. ASCL1 expression is mostly lost in quiescent NSCs, however induced oscillatory *Ascl1* expression in the SGZ of adult mice promotes activation of quiescent NSCs to generate neurons (Sueda et al., 2019). Accordingly, light-induced *Ascl1* oscillations, but not sustained expression, more efficiently activates cultured quiescent NSCs to differentiate (Sueda et al., 2019).

On the other hand, HES1 is expressed at high level in slow dividing boundary regions in the roof and floor plate of mouse spinal cord and was found to oscillate at high level in quiescent NSCs (Baek et al., 2006; Sueda et al., 2019). Deletion of *Hes1* and *Hes1*-related genes affects boundary formation in the embryo and increases neurogenesis. Similarly, in the adult brain it transiently increases neurogenesis followed by NSC depletion, suggesting that *Hes1* is required to maintain NSCs (Baek et al., 2006; Sueda et al., 2019). Moreover, forced expression of sustained *Hes1* has been shown to prevent neural differentiation (Ishibashi et al., 1994; Baek et al., 2006; Sueda et al., 2019), it reduces cell proliferation (Shimojo et al., 2008; Baek et al., 2006) and maintains NSCs in a quiescent state (Sueda et al., 2019). Therefore, high sustained or high oscillatory HES1 expression represses proneural gene expression and inhibits NSC proliferation. Unlike ASCL1 though, it is not clear whether the dynamic expression of HES1 itself, rather than the high level, can drive this phenotype.

The significance of HES1 oscillations has been previously studied *in vivo* during early mouse development. Generation of a mutant *Hes1* mouse carrying a shorter *Hes1* gene leads to dampening of HES1 oscillations while maintaining the HES1 level above the required total amount for normal development. Comparison of these mutant embryos versus wild-type (WT) *Hes1* embryos in a *Hes3/Hes5*-null background showed that in the absence of HES1 oscillations neurogenesis was accelerated whereas the size of the telencephalon was decreased (Ochi et al., 2020). However, the role of HES1 dynamics on the ability of NSCs to enter and/or exit quiescence still remains unknown.

Here, we provide a detailed characterisation of HES1 expression in NSCs as they transit in and out of quiescence *in vitro* and we explore the role of HES1 dynamics during this transition. We find that upon induction of NSC quiescence with BMP4 the total HES1 level does not change while the HES1 oscillations are retained. To assess the role of HES1 dynamics in driving entry or exit from quiescence we first generated a new fluorescent *HES1*^*mScarlet-I/mScarlet-I*^ transgenic mouse line, where the gene encoding for the mSCARLET-I fluorophore has been fused to the endogenous *Hes1* locus, and we established NSC cultures from homozygote (HOM) mice where mSCARLET-I represents total endogenous HES1 expression. We then introduced a sustained mVENUS:HES1 input under a moderate-strength UbC promoter (UbC-mVENUS:HES1) and assessed its effect on endogenous HES1 dynamics and HES1 level. We found that upon expression of UbC-mVENUS:HES1 the endogenous HES1 oscillations were abolished and endogenous HES1 level decreased by x3.5fold. Absolute quantification of the HES1 concentration with fluorescence correlation spectroscopy (FCS) showed that although the total HES1 concentration was increased in UbC-mVENUS:HES1 expressing cells, this was still within the range of HES1 concentrations found in cells not expressing the sustained input. Interestingly, under these conditions NSCs continue to proliferate and they can be induced into quiescence however, exit from quiescence is impeded. We therefore provide evidence that oscillatory HES1 dynamics are required specifically for reactivation of NSCs from quiescence.

## Results

### 1. HES1 oscillations persist through quiescence and reactivation, and are of better quality in quiescent NSCs

To monitor the HES1 dynamics in active and quiescent NSCs, we first established NSC lines from the telencephalic lateral ganglionic eminence (LGE) of E13.5 mouse embryos (Fig.1A). The embryos were derived from HES1 reporter mice where the firefly Luciferase 2 (LUC2) cDNA had been inserted in-frame upstream of the *Hes1* gene in a bacterial artificial chromosome (BAC), to generate a LUC2-HES1 fusion protein that mimics the endogenous HES1 expression (Imayoshi et al., 2013) (Fig.1A). Embryonic LGEs were chosen as their descendants contribute to the pool of adult NSCs that reside in the ventral aspect of the subventricular zone (SVZ) (Young et al., 2007).

**Figure 1:**
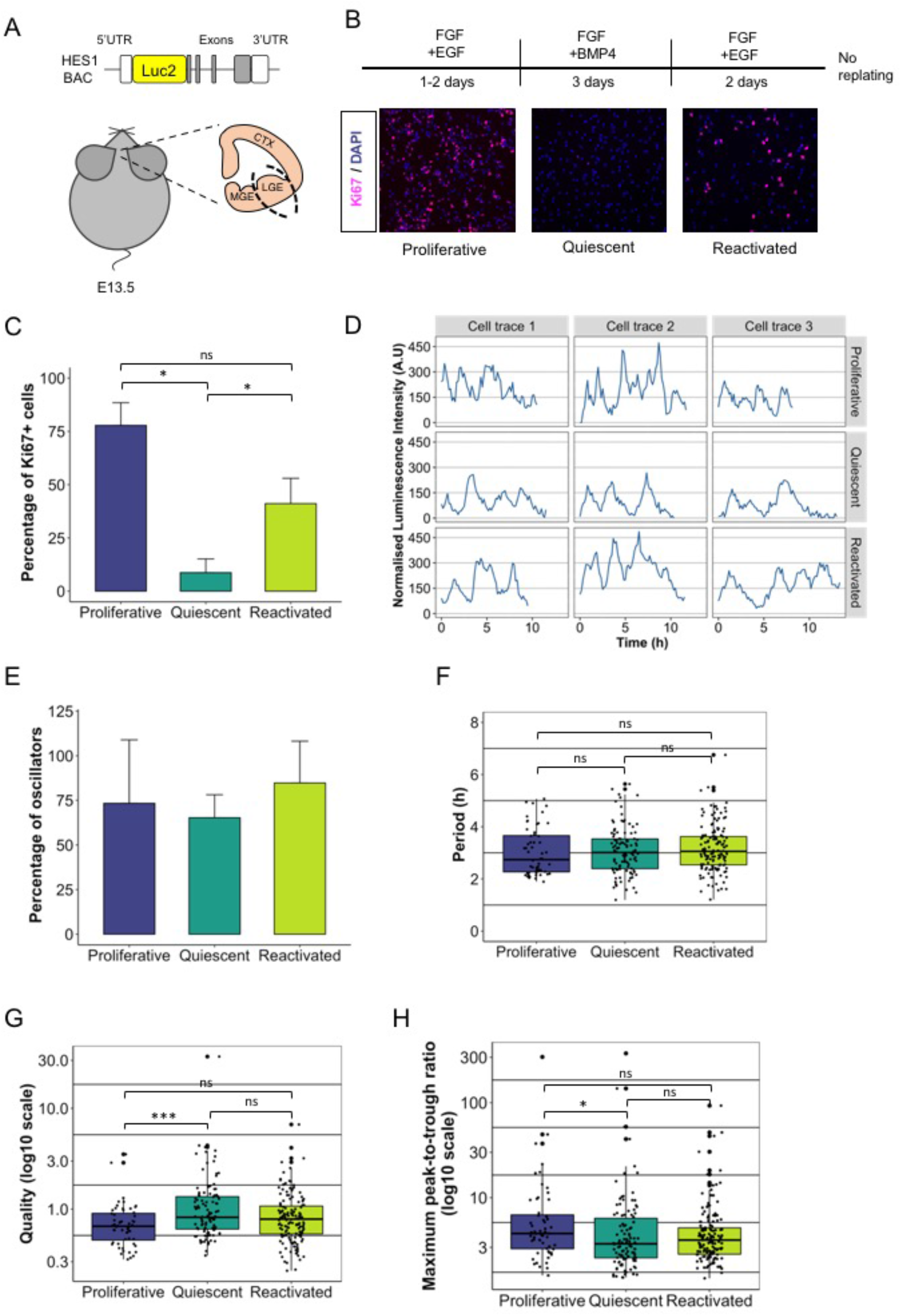
Characteristics of HES1 dynamic expression in proliferative, quiescent and reactivated conditions. A) Schematic structure of LUC2:HES1 BAC used to generate transgenic mice. NSC lines were established from the LGE region of E13.5 LUC2:HES1 embryos (CTX=cortex, LGE=lateral ganglionic eminence, MGE=medial ganglionic eminence). B) Experimental design for induction of quiescence and reactivation (upper panel). Example images from the analysis of cell proliferation by Ki67 staining at proliferative, quiescent and reactivated conditions. Cells were counterstained with DAPI (bottom panels). C) Percentage of Ki67 positive cells in proliferative, quiescent and reactivated conditions (error bars represent standard deviation, n=3 biological experiments, One-way ANOVA with Tukey’s multiple comparison test, *p<0.05, ns=not significant). D) Example cell traces of E13.5 LUC2:HES1 NSCs cultured in proliferative, quiescent and reactivated conditions. E) Percentage of oscillators in proliferative, quiescent and reactivated conditions (Error bars represent standard deviation, number of 10h cell traces analysed: 75 proliferative from n=8, 171 quiescent from n=4, 182 reactivated from n=5 biological experiments). F-H) Box plots representing analysis of period of oscillations (F), quality of oscillations (G) and maximum peak-to-trough ratio per oscillatory trace (H) (dots represent individual oscillatory cell traces, black horizontal lines represent median, number of 10h oscillatory cell traces analysed: 56 proliferative from n=7, 110 quiescent from n=4, 149 reactivated from n=5 biological experiments, Kruskal-Wallis test with Dunn’s multiple comparison test, *p<0.05, ***p<0.001, ns=not significant)

E13.5 LUC2-HES1 LGEs were cultured in the presence of epidermal growth factor (EGF) and basic fibroblast growth factor (FGF) to promote their proliferation, as indicated by the expression of the cell proliferation marker Ki67 (Fig.1B-C), and they uniformly expressed the stem cells markers SOX2 and PAX6 suggesting that they retain their stem cell characteristics in culture (Suppl. Fig.1A). To induce quiescence, EGF was replaced by BMP4 for 3 days upon which the NSCs lost their Ki67 expression and they were cell-cycle arrested (Fig.1B-C). It has been previously shown that addition of BMP4 in the presence of FGF activates a quiescence gene expression programme (Martynoga et al., 2013). The main mediators of BMP signaling were found to be ID proteins (inhibitors of differentiation), and not SMADS, although *Id* genes were not considered to be the main mechanism by which quiescence was established (Martynoga et al., 2013). Accordingly, we find in our cultures that upon addition of BMP4 all *Id* genes (*Id1*-*4*) are upregulated with *Id4* upregulated the most (Suppl. Fig.1B). To reactivate the cells, BMP4 was replaced by EGF, enabling the cells to re-enter the cell-cycle and start proliferating again (Fig.1B-C) (Martynoga et al., 2013). To ensure that upon reactivation we are not promoting the proliferation of the minority of cells that failed to enter quiescence, we reactivated the cells without replating and only by replacing the culture media (Fig.1B).

We next performed bioluminescent single-cell live-imaging of NSCs cultured in either proliferative, quiescent or reactivated conditions. HES1 was expressed and expression was dynamic under all three conditions (Fig.1D). Proliferating NSCs are characterised by increased cell motility which restricts the amount of time individual cells can be tracked for before they exit the field of view, whereas quiescent cells are less motile and therefore can be tracked for longer (Suppl. Fig.1C-D). To therefore avoid any bias introduced by the length of tracking we truncated all cell traces to have a maximum length of 10h, to match the average cell trace length in proliferating conditions (Suppl. Fig.1D). For example, traces that were 30h long they were split into 3 independent 10h traces, so that no information was lost.

We first sought to determine whether the HES1 dynamic expression was oscillatory by using a previously described GP-based method (see Materials and Methods) that can classify cell traces as periodic or not, with statistical significance (Phillips et al., 2017). Our analysis revealed that across all different conditions ∼70% of cell traces were oscillatory (Fig.1E). HES1 was found to oscillate in proliferative conditions with a median period of 2,7h, similarly to what has been previously described (Imayoshi et al., 2013), whereas in quiescent and reactivated conditions the HES1 median period was slightly increased by ∼20min, although it was not statistically significant (Fig.1F).

We also assessed and compared the quality of oscillations, which is the ratio of the frequency of oscillations to the timescale of damping (Phillips et al., 2017). Oscillations with high quality better match a sine wave whereas low quality oscillations tend to have higher peak-to-peak variability. We found that HES1 oscillations in quiescent cells are of better quality (Fig.1G). Finally, we compared the amplitude of HES1 oscillations and found that proliferative cells tend to have a higher peak-to-trough ratio (Fig.1H). Oscillations in reactivated cells showed a mixed ‘phenotype’ with intermediate levels of quality and amplitude, indicative of their tendency to return to a proliferative state (Fig.1G-H). Overall, our results show that HES1 continues to oscillate in quiescent and reactivated NSCs, similarly to proliferative conditions, with HES1 oscillations being of better quality in quiescent NSCs.

### 2. HES1 level does not increase in BMP4 induced quiescent NSCs

High HES1 expression has been reported to associate with slow-dividing or quiescent cells whereas enforced high-sustained expression prevents cell proliferation (Baek et al., 2006; Shimojo et al., 2008; Sueda et al., 2019) suggesting that high HES1 is potentially required for quiescence to be established. To assess how HES1 level is affected upon induction of quiescence with BMP4 in our system, we performed bioluminescent live-imaging of LUC2-HES1 NSCs as they transition from proliferation into quiescence. We began imaging NSCs in proliferating conditions and then replaced the media to induce quiescence while continuing to image. Following media replacement, the cells were imaged for at least three more days, by the end of which the vast majority of the cells have become quiescent (Fig.1B-C). Individual cells expressing luminescence were tracked before and after induction of quiescence to record HES1 levels in three independent experiments (Fig.2A). To assess the HES1 level we first estimated the median luminescence expression from all cell traces at every timepoint and then compared the median expression of all sampling intervals between proliferative and quiescent cells. Our analysis revealed that they were no significant differences in HES1 level between proliferative and quiescent NSCs (Fig.2B). Similar results were obtained when we compared the median luminescent expression per cell trace in proliferative versus quiescent conditions (Suppl. Fig2A). To avoid any bias introduced by the fact that quiescent cells can be tracked for longer, where potentially more low expressing intervals can be captured, we performed additional analysis to overcome this limitation. For each replicate experiment, we calculated the median luminescence expression across both proliferative and quiescent conditions, which was estimated based on the median HES1 expression at each sampling interval, and we used it as a threshold (Fig.2A,C). We then explored for how long the HES1 level was detected above the threshold, continuously. That is, for every trace we would estimate continuous blocks of time intervals during which HES1 was expressed above the threshold. Each block of time interval represents an oscillatory peak and it was treated as an independent component. Blocks of intervals of 30min or less were excluded from any downstream analysis as they were too short to be considered oscillatory peaks. Analysis of the continuous time spent above threshold between proliferative and quiescent conditions showed variable behaviour but not statistically significant differences (Fig.2D). We also measured the area under the curve for each peak that had crossed the threshold, to assess how much HES1 protein was expressed (Fig.2E). Our results showed again no consistent requirements for HES1 level to increase or decrease in order for the cells to enter quiescence (Fig.2E). Finally, we compared the maximum fold change of each peak over the equivalent threshold value, where we found again no statistically significant differences (Fig.2F). Together these findings show that upon induction of quiescence with BMP4 the HES1 level does not change.

**Figure 2:**
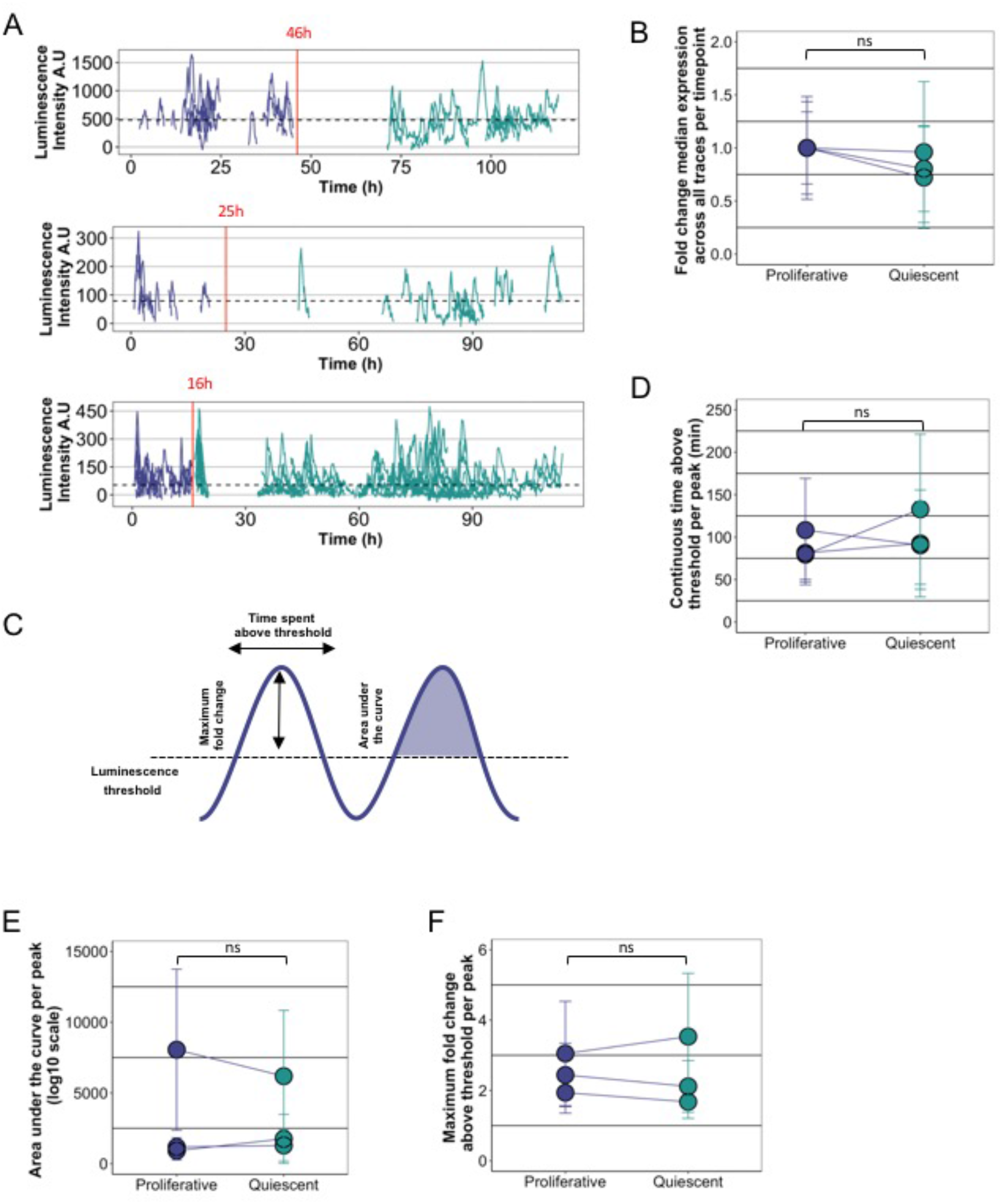
HES1 level does not change as NSCs transition from proliferation into quiescence. A) Graphs showing luminescence expression of LUC2:HES1 NSCs as they transition from proliferation into quiescence from 3 independent experiments. Red vertical lines indicate the time at which proliferation medium was replaced by quiescence medium and horizontal dotted lines represent the luminescence thresholds defined as the median luminescence expression across proliferative and quiescence conditions per experiment, estimated by the median luminescence expression at each sampling interval. B) Graph showing the relative fold change of median luminescence expression between proliferative and quiescent conditions per experiment. Luminescence expression was estimated by the median luminescence expression at each sampling interval across proliferative and quiescent cell traces (error bars represent standard deviation, two-tailed paired t-test, ns=not significant). C) Schematic showing the different parameters tested for each peak from all cell traces. These include the maximum fold change of luminescence expression per peak relative to the threshold, the continuous time spent above the threshold for every peak and the area under the curve per peak. D-F) Graphs comparing the time spend above threshold per peak (D), the area under the curve (E) and the maximum fold change above threshold per peak (F) between proliferative and quiescent conditions per experiment (circles represent mean values and error bars represent standard deviation, two-tailed paired t-test, ns=not significant)

### 3. Ectopic sustained HES1 expression overrides the endogenous HES1 oscillations

To further gain insight into the functional role of the HES1 dynamics, we next sought to establish a system to manipulate HES1 oscillatory expression. To this end, we first generated a transgenic mouse HES1 reporter line to mark endogenous HES1 expression by knocking-in the mSCARLET-I fluorophore, in frame, at the C-terminus of the endogenous *Hes1* locus (Fig.3A, Suppl. Fig.3A). E13.5 LGEs were further isolated from *HES1*^*mScarlet-I/mScarlet-I*^ mice to establish NSCs lines where mSCARLET-I expression represents the total expression of endogenous HES1 in the cells.

**Figure 3:**
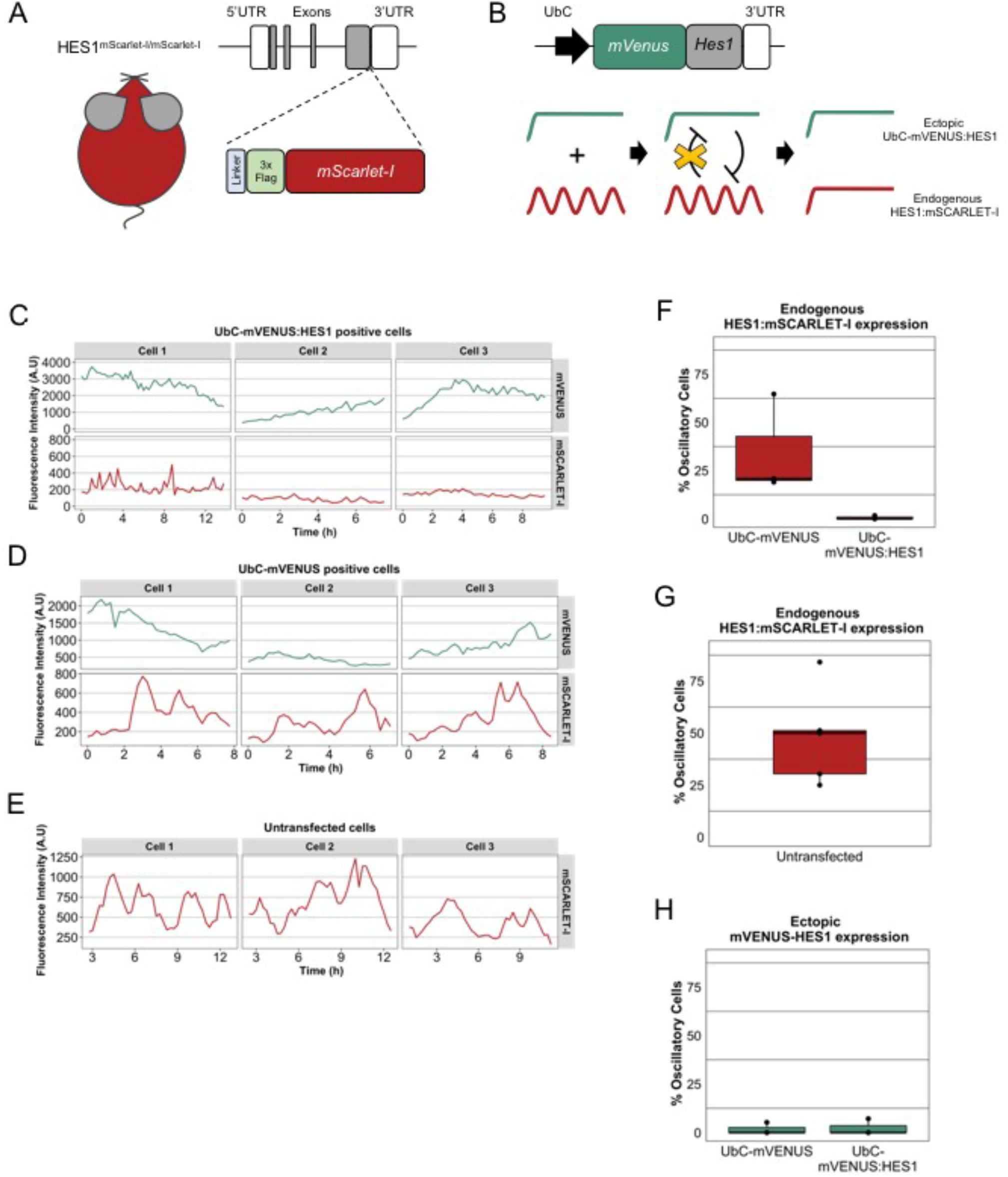
Endogenous HES1 protein does not oscillate in the presence of an ectopic HES1 sustained input. A) Schematic structure of the *Hes1* locus in the *HES1*^*mScarlet-I/mScarlet-I*^ transgenic mice. A DNA sequence encoding for a linker protein, a 3xFlag epitope and the mSACRLET-I protein has been inserted downstream of the last *Hes1* exon and before the 3’UTR. B) Schematic showing the structure of UbC-mVENUS:HES1 reporter and the hypothesis for altering the endogenous HES1:mSCARLET-I oscillatory expression. *mVenus* has been fused to *Hes1* cDNA followed by the *Hes1* 3’UTR and it is expressed under the constitutive UbC promoter. C-E) Example cell traces showing mVENUS and mSCARLET-I fluorescence expression over time in the same cell where mVENUS represents expression of either UbC-mVENUS:HES1 (C) or the control UbC-mVENUS (D) and mSCARLET-I represents endogenous HES1 expression. In untransfected cells only the mSCARLET-I endogenous HES1 expression has been recorded (E). F-G) Box plots showing the percentage of cells that have oscillatory endogenous HES1:mSCARLET-I expression in the presence of ectopic UbC-mVENUS or UbC-mVENUS:HES1 (F) or in untransfected cells (G). H) Box plot showing the percentage of cells where UbC-mVENUS or UbC-mVENUS:HES1 exhibit oscillatory expression

To manipulate the endogenous HES1 dynamics we introduced into the *HES1*^*mScarlet-I/mScarlet-I*^ NSCs an mVENUS:HES1 reporter driven by the ubiquitously expressed and moderate-strength UbC promoter. The notion behind this is that ectopic expression of a sustained HES1 input (i.e. UbC-mVENUS:HES1) will continuously suppress the endogenous HES1 expression (due to HES1 autorepression) while the endogenous HES1 protein will not be able to bind and inhibit the UbC-mVENUS:HES1 reporter, as it lacks the HES1 binding sites (Fig.3B). Therefore, we hypothesised that ectopic sustained HES1 expression will override the endogenous HES1 oscillations.

To assess whether this is the case, we performed dual-colour fluorescence live-imaging of *HES1*^*mScarlet-I/mScarlet-I*^ NSCs expressing the UbC-mVENUS:HES1 reporter, to record the endogenous HES1 dynamics in the presence of a HES1 sustained input. We then compared the results against *HES1*^*mScarlet-I/mScarlet-I*^ NSCs expressing a control UbC-mVENUS reporter or no reporter at all. We found that HES1:mSCARLET-I endogenous expression continued to be dynamic in untransfected cells or cells expressing the control UbC-mVENUS whereas the oscillations were completely abolished in cells expressing the UbC-mVENUS:HES1 (Fig.3C-E). Oscillatory analysis of the traces using the GP-based method previously described confirmed that endogenous HES1:mSCARLET remains oscillatory in untransfected and control UbC-mVENUS trasnfected cells whereas almost no oscillations were found in cells expressing UbC-mVENUS:HES1 (Fig.3F-G). Finally, as expected, neither the ectopic sustained UbC-mVENUS:HES1 expression nor the control UbC-mVENUS expression were found to be oscillatory as expression of these reporters is driven by the constitutive UbC promoter (Fig.3H). We therefore conclude that the ectopic sustained HES1 expression overrides the endogenous HES1 oscillations and can be used as a means to manipulate the endogenous HES1 dynamics.

### 4. Sustained HES1 expression impedes reactivation of quiescent NSCs

Having established that we can override the endogenous oscillations by introducing ectopically a sustained HES1 input, we next sought to determine the effect of altering the HES1 dynamics on the ability of NSCs to proliferate as well as to enter and exit quiescence. To address that we employed a similar method as before whereby the UbC-mVENUS:HES1 reporter was transfected into WT NSCs to create a mosaic population of UbC-mVENUS:HES1 positive and negative cells. We then cultured the cells either in proliferative conditions or we induced quiescence for 3 days, followed by reactivation into proliferating conditions again for 2 days. At each condition we fixed the cells and performed immunofluorescent staining against the Ki67 proliferation marker and mVENUS (Fig.4A). As a control, we applied the same process to NSCs transfected with the UbC-mVENUS reporter (Fig.4B).

**Figure 4:**
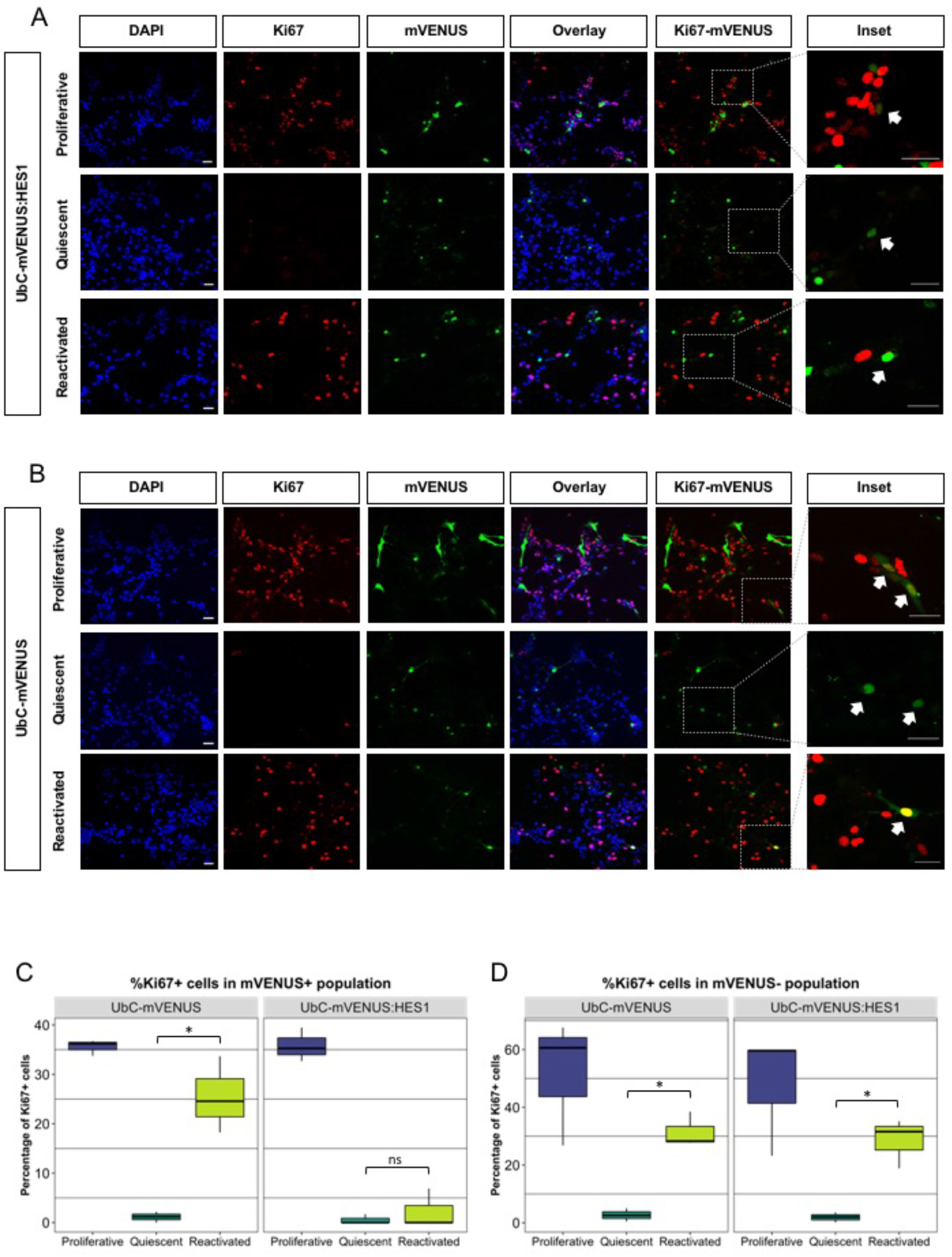
Loss of HES1 oscillations in NSCs impedes reactivation from quiescence. A-B) Immunofluorescence staining for the proliferation marker Ki67 and mVENUS. E13.5 NSCs were transfected either with the UbC-mVENUS:HES1 (A) or the control UbC-mVENUS reporter (B). Cells were fixed and stained in proliferative, quiescent or reactivated culture conditions. Cells were also counterstained with DAPI. The insets depict the dotted square areas magnified. White arrows in the insets in proliferative conditions in (A) and (B) mark cells that stain for both Ki67 and mVENUS whereas in quiescent conditions in (A) and (B) white arrows mark mVENUS positive cells that are negative for Ki67. In reactivated conditions in (A) white arrow marks a cell that stains positive only for mVENUS but it is negative for Ki67 whereas in (B) the arrow marks a cell that stains positive for both mVENUS and Ki67 (scale bars=30μm). C-D) Box plots showing the percentage of Ki67 positive cells in UbC-mVENUS or UbC-mVENUS:HES1 negative (C) and positive cells (D) in proliferative, quiescent and reactivated conditions (black horizontal lines represent median, n=3 biological experiments, two-tailed paired t-test, *p<0.05, Wilcoxon test (for comparison in mVENUS positive in UbC:mVENUS:HES1 transfected cells), ns = not significant).

We estimated the percentage of Ki67 positive cells at each condition and within the mVENUS positive and mVENUS negative populations. We found that within the mVENUS positive population (i.e. transfected cells) in proliferative conditions UbC-mVENUS:HES1 expressing cells were proliferating at the same rate as the control UbC-mVENUS expressing cells, suggesting that switching from oscillatory to stable HES1 expression does not prevent cells from cycling (Fig.4C). Equally, upon induction of quiescence, almost all UbC-mVENUS:HES1 expressing cells lost Ki67 expression, similar to control, indicating that dynamic expression of HES1 is not required for cells to enter quiescence (Fig.4C). Interestingly though, when UbC-mVENUS:HES1 expressing quiescent cells were exposed to proliferating media they did not resume proliferation (Fig.4C). This impediment in reactivation was only present in UbC-mVENUS:HES1 expressing cells but not in their control counterparts. Accordingly, we found that within the mVENUS negative population (i.e. untransfected cells), in both the control UbC-mVENUS and the UbC-mVENUS:HES1 treated cultures, the NSCs were able to proliferate when cultured in proliferating conditions and they would undergo quiescence when exposed to BMP4 (loss of Ki67 expression) whereas upon reactivation they all resumed proliferation capacity (Fig.4D). These findings suggest that oscillatory HES1 expression is required specifically for cells to be reactivated from quiescence.

### 5. Total HES1 level remains within physiological level range in cells expressing sustained HES1

To further determine whether the failure of UbC-mVENUS:HES1 expressing NSCs to reactivate is due to the change in HES1 dynamic expression rather than the HES1 protein level, it is imperative to assess how the total HES1 level is affected in the presence of the ectopic sustained HES1 input.

We first looked at how the endogenous HES1 level was affected in the presence of UbC-mVENUS:HES1 or control UbC-mVENUS reporter. From the dual-colour fluorescence imaging movies that we previously generated (Fig.3C-E) we estimated the mean endogenous HES1:mSCARLET-I fluorescence intensity per cell trace in the presence or absence of an ectopic reporter. In particular, we found that in control UbC-mVENUS expressing cells the endogenous HES1 expression was not significantly altered when compared to untransfected cells however, in the presence of ectopic sustained HES1 the endogenous HES1 level was decreased by a x3.5fold (Fig.5A-B). These results confirm our expectation that ectopic sustained HES1 would repress the endogenous HES1 expression.

This decrease in the endogenous HES1 levels however does not inform on how the total HES1 level is affected in UbC-mVENUS:HES1 expressing cells, since we cannot estimate the amount of the mVENUS:HES1 protein that we introduce based on fluorescence intensity. To this end, we employed fluorescence correlation spectroscopy (FCS) to perform absolute quantification of HES1 protein molecules. FCS analysis measures the fluctuations in concentration of fluorescent particles in a minute volume (also known as confocal volume) and temporally autocorrelates the recorded intensity signal to infer absolute particle numbers. We established mosaic cultures of *HES1*^*mScarlet-I/mScarleti-I*^ NSCs expressing the sustained UbC-mVENUS:HES1 and we performed a dual-colour FCS for mSCARLET-I and mVENUS (Fig. 5Ci). In particular, absolute quantification of mSCARLET-I fluorescent particles per confocal volume would be a direct estimation of the endogenous HES1 concentration whereas quantification of mVENUS particles would be a direct estimation of HES1 concentration expressed under the UbC promoter.

**Figure 5:**
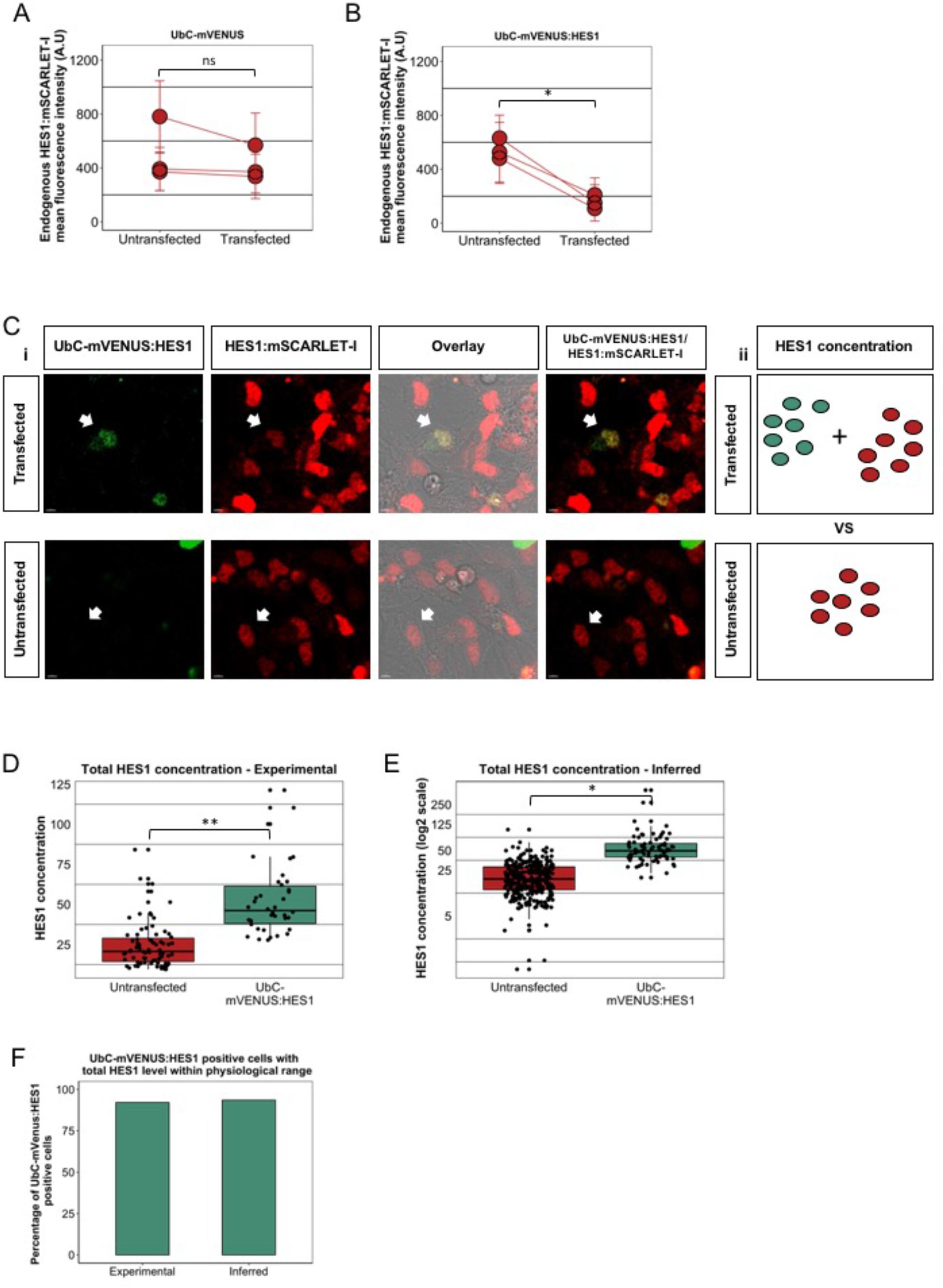
Ectopic sustained HES1 expression does not increase total HES1 level above physiological range. A-B) Graphs showing the endogenous HES1:mSCARLET-I expression level in untransfected vs transfected cells with UbC-mVENUS (A) or UbC-mVENUS:HES1 reporter (B) within the same culture per experiment (circles represent mean values and error bars represent standard deviation, number of cells analysed for A) untransfected=191, transfected=135, n=3 biological experiments, number of cells analysed for B) untransfected=222, transfected=88, n=3 biological experiments, two-tailed paired t-test, *p<0.05, ns=not significant). C) Example pictures of E13.5 HES1:mSCARLET-I NSCs transfected with UbC-mVENUS:HES1 reporter. White arrows mark transfected (top panels) or untransfected (bottom panels) cells where FCS was performed (i). Schematic showing how total HES1 concentration is compared between transfected and untransfected cells. Green dots represent UbC-mVENUS:HES1 concentration and red dots represent HES1-mSCARLET-I concentration (scale bars=5μm). D-E) Box plots showing total HES1 concentration in untransfected and UbC-mVENUS:HES1 transfected E13.5 HES1:mSCARLET-I NSCs. In the reporter transfected cells, the total HES1 concentration is the sum of UbC-mVENUS:HES1 concentration and HES1:mSCARLET-I concentration. In D) all concentrations have been estimated experimentally by FCS (apart from the HES1:mSCARLET-I concentration in transfected cells which was inferred). In E) all concentrations have been inferred (black horizontal lines represent median, number of cells analysed in D) untransfected= 75, transfected=38, n=3 biological replicates, in E) untransfected=316, transfected=77, n=3 biological experiments, two-tailed paired t-test, *p<0.05, **p<0.01). F) Bar graph showing the percentage of E13.5 HES1:mSCARLET-I NSCs transfected with UbC-mVENUS:HES1 that have a total HES1 concentration (either experimentally estimated or inferred) within the range of total HES1 concentration values in their untransfected counterparts.

To estimate the total concentration of HES1, the number of mVENUS and mSCARLET-I HES1 molecules per confocal volume were combined in UbC-mVENUS:HES1 expressing cells and compared against the HES1:mSCARLET-I molecules in untransfected cells (Fig. 5Cii). However, we found that in cells expressing the UbC-mVENUS:HES1 the endogenous HES1 (in the vast majority of the cells) was so dramatically decreased that the amount of HES1 molecules could not be accurately measured by FCS. To therefore ensure that we are not underestimating the endogenous HES1 expression in those cells, by assuming zero contribution when an accurate measurement cannot be obtained, we decided instead to infer the HES1 concentration from the mSCARLET-I fluorescence intensity. For this purpose we constructed standard curves of mSCARLET-I and mVENUS fluorescence intensity versus mSCARLET-I concentration (in unstransfected cells) and mVENUS concentration (in UbC-mVENUS:HES1 expressing cells) respectively in the same cell, from 3 independent experiments (Suppl. Fig.4A). In each case we found a strong correlation of 0.8 or greater, suggesting that the estimation of HES1 concentration is representative of the amount of the HES1 protein expressed in each cell.

We performed two types of analyses in order to estimate the total amount of HES1 protein. One analysis was based on the experimental values obtained by FCS (called experimental analysis), with the exception of the endogenous HES1:mSCARLET-I particles in the UbC-mVENUS:HES1 expressing cells which had to be inferred from the fluorescence intensity. For the second analysis we inferred all HES1 concentration based on mSCARLET-I and mVENUS fluorescence intensity from cells where no FCS was performed (called inferred analysis). The aim of the latter analysis was to increase the number of cells tested in order to get a better representation of the HES1 level heterogeneity in the culture.

Overall, we found that total HES1 levels in UbC-mVENUS:HES1 expressing cells increased by either x2.1fold (experimental analysis) or x2.9fold (inferred analysis) (Fig.5D-E). However, over 90% of the UbC-mVENUS:HES1 expressing cells had a total HES1 concentration still within the range of HES1 concentrations recorded in untransfected cells (which we consider to represent the physiological range of HES1 level) (Fig.5F). These results suggest that despite the overall increase in total HES1 level, cells expressing a HES1 sustained input operate within normal HES1 level. We also calculated the contribution of endogenous HES1-mSCARLET-I and ectopic mVENUS-HES1 concentration in the total HES1 level of UbC-mVENUS:HES1 expressing cells (in both the experimental and inferred analysis) and found an approximate x4fold decrease in endogenous HES1 levels compared to untransfected cells (Suppl. Fig.4B-C), similar to what we found before based on the fluorescence intensity comparison (Fig.5B). Altogether these findings reveal that ectopic expression of sustained HES1 under a weak ubiquitous promoter does not increase total HES1 level above normal values, revealing that the impediment of reactivation in UbC-mVENUS:HES1 expressing cells is due to the change in HES1 dynamics and not levels.

## Discussion

Here we have explored the role of HES1 ultradian oscillations in controlling the entry and exit of NSCs from quiescence *in vitro*. We showed that by altering the HES1 dynamic expression from oscillatory to sustained, while keeping the total HES1 level within physiological range, reactivation from quiescence is impeded while NSC proliferation and entry into quiescence is not affected.

Our findings highlight the importance of ultradian gene expression dynamics regulating cell fate, a concept that has been evolving over the last several years (Isomura and Kageyama, 2014) as more and more evidence linking the temporal regulation of ultradian oscillations with distinct cellular outcomes emerges (Imayoshi et al., 2013; Shimojo et al., 2016; Santos et al., 2007; Purvis et al., 2012). For example, the Notch ligand Delta-like1 (DLL1) and the transcriptional repressors HES1 and HES7, they all exhibit oscillatory expression in the mouse presomitic mesoderm. Elongation or shortening of the of the *Dll1* gene dampens the DLL1 oscillations, followed by a dampening of HES1 and HES7 oscillations, which lead to severe fusion of somites and their derivatives (Shimojo et al., 2016). Accordingly, the tumour suppressor protein P53 exhibits oscillatory expression (with a period of ∼5.5h) upon exposure to γ-irradiation (Lev Bar-Or et al., 2000; Lahav et al., 2004), which allows for cells to recover from DNA damage and survive, while induction of sustained P53 signaling leads to senescence (Purvis et al., 2012). Therefore, understanding how gene expression dynamics dictate cell fate is essential for being able to manipulate cell-state transitions at will.

High sustained HES1 expression has been previously shown to associate with slow-diving cells *in vivo* or prevent NSC proliferation both *in vitro* and *in vivo*, while quiescent NSCs were found to express high oscillatory HES1 (Baek et al., 2006; Shimojo et al., 2008; Sueda et al., 2019). In all cases though, HES1 is expressed at a high level which does not allow to dissect out how each factor (HES1 level vs HES1 dynamic expression) contributes to this phenotype. Here we set out to explore the role of HES1 expression dynamics in the context of NSC quiescence. We first investigated how is the HES1 dynamic expression affected when proliferative NSCs are induced in and out of quiescence *in vitro* with the addition and subsequent removal of BMP4. Similarly to previously published data (Sueda et al., 2019) we found that HES1 oscillatory expression persists throughout quiescence and reactivation. We did not find significant differences in the period of oscillations but we did find that oscillations in quiescent cells were of better quality. This could be due to the fact that quiescent cells are less motile and interact with fewer neighbours, therefore Notch signaling in these cells is less interrupted by intercellular Notch lateral inhibition (Kageyama et al., 2019).

With regards to HES1 levels, we did not find an increase in the HES1 level overall or a consistent pattern of HES1 oscillatory peaks reaching higher levels or being wider compared to proliferative conditions. This is in contrast to what had been previously reported where HES1 level was found to increase upon induction of quiescence in embryonic NSCs *in vitro* (Sueda et al., 2019). However, it is possible that the use of different BMP ligands to establish quiescence or the use of the same ligand at a different concentration to differentially affect the level of HES1. In our cultures we find that BMP4 addition induces expression of all *Id* genes but *Id4* is the most upregulated. ID proteins have an HLH binding domain but they cannot bind onto DNA (Ruzinova and Benezra, 2003) and they act primarily by interfering with the transcriptional function of proneural proteins though sequestration of E proteins, which heterodimerize with bHLH transcription factors like ASCL1 (Langlands et al., 1997; Vinals et al., 2004; Sharma et al., 2015). All ID proteins have been reported to interact with HES1 (Jogi et al., 2002). In particular ID2 has been found to release the negative autoregulation of HES1 and increase HES1 level without interfering though with HES1 repression on its downstream targets (Bai et al., 2007). However, ID4, appears to have distinct functions to the rest of IDs (Patel et al., 2015). It has been shown that ID4 can heterodimerise with ID1, ID2 and ID3 and inhibit their biological activity (Sharma et al., 2015) and in this way retain the HES1 oscillations by preventing HES1 to interact with the other IDs (Boareto et al., 2017). It is therefore possible that during establishment of quiescence in response to BMP signaling, differential expression of ID proteins can differentially affect HES1 level, with ID2 for example increasing HES1 level and ID4 inhibiting the function of ID2, thus preventing upregulation of *Hes1*.

To assess the role of HES1 dynamics in controlling the entry and exit of NSCs from quiescence we first generated a new HES1:mSCARLET transgenic mouse line, where both alleles of the *Hes1* locus were tagged with the mSCARLET-I fluorophore, to monitor HES1 endogenous expression. By introducing ectopically a sustained HES1 input, under the control of a moderate-strength UbC promoter, we found that we can override the endogenous HES1 oscillations without increasing the total HES1 level above the physiological range. Under these conditions proliferation of NSCs did not seem to be affected. This is line with what has been previously reported where NSCs derived from HES1 mutant mice, with dampened HES1 oscillations, showed no differences in proliferation and neuronal differentiation compared to WT NSCs *in vitro* (Ochi et al., 2020). Accordingly, mathematical modelling of the Notch/HES regulatory module in NSCs predicted that loss of HES1 oscillations are dispensable for low proneural factor activity that is required for NSC proliferation (Boareto et al., 2017).

We then assessed how loss of HES1 oscillations affects entry into quiescence and reactivation. Upon induction of quiescence, NSCs expressing a sustained HES1 input stopped proliferating, similar to control cells, however, upon reactivation they failed to re-renter the cell-cycle. We cannot conclude though whether this is a complete block from exiting quiescence or a delayed reactivation. Varying depths of quiescence, accounting for differences in responsiveness to reactivation, have been previously described in adult NSCs but also other systems such as muscle stem cells (MuSCs) and haematopoietic stem cells (HSCs) (Llorens-Bobadilla et al., 2015; Rodgers et al., 2014; Laurenti et al., 2015). Thus, it is possible that sustained HES1 expression alters the depth of quiescence in NSCs. But how can this be achieved? It has been previously reported that changes in the pattern of expression of a transcription factor can differentially affect expression of downstream targets (Ashall et al., 2009; Lane et al., 2017; Hao and O’Shea, 2011; Hansen and O’Shea, 2013; Chen et al., 2020) For example, it has been recently shown using elegant optogenetic tools, that downstream targets of the Crz1 transcription factor in yeast have higher gene expression in response to pulsatile input whereas others responded better to continuous inputs (Chen et al., 2020). Similarly, downstream targets of HES1 may respond differently to sustained versus oscillatory HES1 expression. In the context of our study, HES1 sustained expression may directly or indirectly affect targets involved in the transition from a quiescent to a proliferating state. Such targets may include members of the Rb-E2F pathway which has been shown to function as bistable switch that controls entry into the cell-cycle (Yao et al., 2008). Alternatively, sustained HES1 expression may act as a ‘sponge’ of ID proteins which are overexpressed in response to BMP4. Dimerization of ID proteins with E proteins have been found to increase the half-life of IDs (Bounpheng et al., 1999; Lingbeck et al., 2005) thus, it is possible that sustained HES1 expression may ‘trap’ ID proteins and protect them from degradation and therefore retain them in the cells for longer even after removal of the BMP4 stimulus.

We therefore provide evidence that the HES1 dynamics are required for NSCs to exit quiescence *in vitro*, suggesting that oscillations are needed for cells to transition from one state to another rather than just maintaining a progenitor state. This revised view on the functional significance of HES1 is consistent with previous findings showing that when Her6 (orthologue of mouse HES1 in zebrafish) oscillations are diminished due to the increase of noise, cells are unable to transition from progenitor state to differentiation in zebrafish (Soto et al., 2020) whereas the quality and occurrence of HES5 oscillations in mouse spinal cord increases as cells approach the transition to differentiation (Manning et al., 2019). The significance of HES1 oscillations in controlling the state of quiescence may also have implications in tissue regeneration and in cancer where quiescent cancer stem cells are considered to drive cancer progression and metastasis (Chen et al., 2016; Lee et al., 2020; Phan and Croucher, 2020). Overall, our findings highlight the importance of oscillatory expression in controlling cell state transitions, in contrast to the prevailing view that they are only needed to maintain a progenitor state.

## Acknowledgements

The authors wish to thank Prof. Steve Pollard and Prof. Fiona Doetsch for advice and discussions, Dr Hayley Bennett and Maj Simonsen Jackson for their work on generating the *HES1*^*mScarlet-I/mScarlet-I*^ mouse transgenic line, Dr Cerys Manning and Dr James Bagnall for help with FCS analysis, MMath Joshua Burton for assistance with the statistical analysis, Dr Michael Lie-A-Ling for his help with subcloning the pRRL-UbC-mVenus:Hes1 construct and all members of the Papalopulu lab for comments on the work. The authors would also like to thank the Biological Services Facility, the Bioimaging Facility (especially Peter March and David Spiller for their help with imaging), the Genome Editing Unit and the Genomic Technologies Core Facility of the University of Manchester for technical support. The Bioimaging Facility microscopes used in this study were purchased with grants from BBSRC, Wellcome and the University of Manchester Strategic Fund. This work was supported by a Sir Henry Wellcome Trust Fellowship to E.M (201380/Z/16/Z) and a Wellcome Trust Senior Research Fellowship to NP (090868/Z/09/Z)

## Author contributions

E.M. and N.P. conceived the study, E.M. performed experiments and data analysis, N.S. generated the control UbC-mVENUS construct and assisted with experiments, V.B. refactored MATLAB code for the detection of oscillations, period, quality and amplitude analysis in bioluminescent and fluorescent single cell traces and assisted with FCS data analysis, J.D performed reactivation experiments, immunofluorescent staining and data analysis for Fig.1B-C, A.A. designed and supervised the project for the generation of the *HES1*^*mScarlet-I/mScarlet-I*^ mouse transgenic line, N.P supervised the study. E.M co-wrote the manuscript with N.P and input from N.S. and V.B.. All authors gave final approval for publication.

## Methods

### Cell Culture

Primary NSCs were isolated from dissected LGEs of E13.5 embryos from LUC2-HES1 BAC (RRID:IMSR_RBRC06013) reporter mice or *HES1*^*mScarlet-I/mScarlet-I*^ mice and cultured in complete NS media as previously described (Pollard, 2013) supplemented with 10ng/ml FGF-2 (PeproTech), 10ng/ml EGF2 (PeproTech) and 2μg/ml laminin (Sigma) (proliferation medium). LUC2-HES1 BAC mice were obtained by the RIKEN BRC through the National Bio-Resource Project of the MEXT, Japan (Imayoshi et al., 2013). For induction of quiescence 22K-32K cells per square centimetre were plated in proliferating medium and 24h later (or when cells reached 60-70% confluency) they were washed twice with 1xPBS while attached, and the medium was replaced with complete NS media supplemented with 20ng/ml FGF-2 and 50ng/ml BMP4 (PeproTech) (quiescence medium). To reactivate the cells, following at least 3 days of culture in quiescent medium, the cells were washed twice with 1xPBS while attached, and the medium was replaced with proliferating medium without replating the cells.

### Cell transfection and qPCR

Primary NSCs were transfected with Lipofectamine 3000 (Invitrogen) according to manufacturers’ instructions. 22K-32K cells per square centimetre were plated in proliferating medium and 24h later (or when cells reached 60-70% confluency) the cells were transfected with 80ng plasmid per square centimetre. To generate the pRRL-UbC-mVenus:Hes1 plasmid, the UbC-mVenus:Hes1 sequence was subcloned into a new lentiviral transfer vector previously described (Seppen et al., 2002; Gilham et al., 2010). Briefly, the UbC-mVenus:Hes1 sequence was released from the pLNT UbC-mVenus:Hes1 plasmid (Phillips et al., 2016) using Pac1 (followed by blunting with T4 DNA polymerase) and KpnI and cloned into the pRRL lenti backbone which was released with Pme1 and KpnI. To generate pLNT-UbC-NuVenus plasmid (nuclear Venus driven by the UbC promoter) the mVenus cDNA was PCR-amplified from the pLNT-UbC-mVenus lenti backbone plasmid (Bagnall et al., 2015) using a forward primer containing SV40 NLS and then cloned back to the same vector.

For qPCR analysis RNA was extracted from E13.5 LUC2:HES1 NSCs cultured in proliferative (1-2days), quiescent (3d) or reactivated (2d) conditions using the RNeasy kit (Qiagen) as per manufacturers’ instructions. cDNA was prepared using Superscript III (Invitrogen) as per manufacturers’ instructions and qPCR for *Id1, Id2, Id3, Id4* and *GAPDH* was performed with SYBR green (Applied Biosystems) using the primers listed in Supplementary Table 1.

### Generation of HES1^mScarlet-I/mScarlet-I^ transgenic mice

We used the EASI-CRISPR strategy (Quadros et al., 2017) to generate a C-terminally tagged HES1:mSCARLET-I mouse. Two sgRNA targeting the STOP codon of the *Hes1* gene were selected using the Sanger WTSI website http://www.sanger.ac.uk/htgt/wge/ (Hodgkins et al., 2015). sgRNA sequences (GTGGCGGAACTGAGAGCCTC-*AGG* and TGAGGCTCTCAGTTCCGCCA-*CGG*) were purchased as crRNA oligos, which were annealed with tracrRNA (both oligos supplied by Integrated DNA Technologies) in sterile, RNase free injection buffer (TrisHCl 1mM, pH 7.5, EDTA 0.1mM) by combining 2.5 μg crRNA with 5 μg tracrRNA, heating to 95°C, and slowly cooling to room temperature. To generate the long single strand DNA donor repair template, a homology flanked flexible linker-3xFLAG-mScarletI DNA sequence was cloned and used as a template in an initial PCR reaction with primers Hes1_lssDNA_F catgctcccggccgccatgggaattcggtaccaacagtgggacctcggt and Hes1 lssDNA R caagttcgtttttagtgtccgtcagaagagagaggtgggctagggactttacgggtagcagtggcctgaggctctcacttgtacagctcgtccatgcc. This amplicon appended the universal pGEM_dual_Bio_F primer sequence (catgggaattcggtac) at the 5’ end, and the lssDNA comprising 5’H(96nt)-linker-FLAG-mScarletI-3’H(88nt) was purified as described (Bennett et al., 2020). For embryo microinjection the annealed sgRNA were complexed with EnGen Cas9 protein (New England Biolabs) at room temperature for 10 minutes, before addition of long ssDNA donor (final concentrations; each sgRNA 20 ng/μl, Cas9 protein 40 ng/μl, lssDNA 10 ng/μl). The injection mix was directly microinjected into C57BL6/J (Envigo) zygote pronuclei using standard protocols. Zygotes were cultured overnight and the resulting 2 cell embryos surgically implanted into the oviduct of day 0.5 post-coitum pseudopregnant mice. Potential founder mice were screened by PCR, using primers that flank the homology arms (Geno F ttgcctttctcatccccaac, Geno R gcagtgcatggtcagtcac), used in combination with internal mScarlet-I primers (mScarlet-I R GTCCTCGAAGTTCATCACGC, mScarlet-I F TCCCCTCAGTTCATGTACGG). 1/8 pups (pup #7) was PCR positive for both reactions, and Sanger sequencing confirmed perfect integration of the gene tag. Note pup #5 demonstrated a positive PCR result for one of the two reactions, perhaps indicating an illegitimate repair event (Codner et al., 2018). Pup #7 was bred forward with a WT C57BL6/J mouse. Germline transmission was confirmed through PCR and sequencing and a colony was established. Both HET and HOM mice develop normally. All animal work was performed under UK Home Office project licences (P08B76E2B) according to the conditions of the Animals (Scientific Procedures) Act 1986.

### Immunofluorescence staining

Cells were fixed in 4% PFA for 25min, followed by permeabilisation for 5min with 0.5% Triton-X-100 diluted in PBS. Serum blocking was performed for 20min at RT in 1xWBR (Western Blocking Reagent) (Sigma) solution. Cells were incubated with primary antibodies at 4oC overnight and with secondary antibodies at room temperature for 40min. Coverslips were mounted using ProLong diamond antifade mountant with DAPI (Invitrogen). Primary antibodies were mouse anit-Ki67 (1:200, BD Pharmigen), rabbit a-SOX2 (1:200, Abcam), rabbit a-PAX6(1:300, BioLegend) and rabbit a-GFP (for mVENUS detection) (1:500, Invitrogen). Secondary antibodies were a-rabbit 488 (1:500, Invitrogen) and a-mouse 568 (1:500, Invitrogen). Slides were imaged with a Nikon Eclipse 80i or an LSM 880 Airy Scan Upright and images were analysed with ImageJ.

### Bioluminescence and Fluorescence imaging

For bioluminescence imaging 200K-300K primary NSCs were plated on 35mm glass bottom dishes (Greiner) in medium supplemented with 1mM D-luciferin (Promega) and imaged on an Olympus UK LV200 inverted bioluminescence microscope using a 40x oil objective. Cells were kept at 37oC in 5%CO2 throughout the imaging period. Images were acquired with a 10minute exposure and a 2×2 binning. Imaging was performed either in proliferating conditions (cells had been cultured in proliferation medium for at least 1day) or quiescent conditions (cells had been cultured in quiescence medium for at least 3 days) or reactivation conditions (cells had been reactivated in proliferation medium for at least 2 days). To perform continuous bioluminescence imaging of cells transitioning from proliferating to quiescent conditions, imaging was paused to wash the cells, while the dish was firmly attached to the stage, and replace the proliferation medium with quiescent, supplemented with fresh 1mM D-luciferin.

For dual fluorescence imaging of mVENUSs and mSCARLET-I fluorophores, 200K-300K primary NSCs were plated on 35mm glass bottom dishes and imaged on a Nikon A1-R confocal microscope using a 20x air objective and with sequential scanning to avoid any bleed through from one channel to another. Images were acquired every 15min-20min. Bioluminescent and fluorescent movies were analysed with the Imaris imaging software and individual cells were manually tracked using the ‘Spots’ function. Background traces (i.e. detection of bioluminescence or fluorescence signal from a cell-free area) were collected alongside from the same field of imaging for every movie that was analysed. Prior to tracking, the bioluminescent images were subjected to 3×3 median filter to remove bright spot artefacts from cosmic rays.

### Detection of oscillations using Gaussian processes

For the detection of oscillators we considered only single cell HES1 timeseries of 5.5h length (approximately double the expected period of 2-3h). The bioluminescence/fluorescence HES1 timeseries of each cell trace was normalised per experiment by background subtraction. For this, the average timeseries of 2-4 backgrounds collected per video were smoothed using a nonlinear polynomial (order 2) to account for minor fluctuations due to detection noise. The smoothed background timeseries was then subtracted from the bioluminescence/fluorescence timeseries corresponding to each cell at each available time point. This produced pre-processed timeseries of HES1 in both bioluminescence and fluorescence experiments.

To identify statistically significant HES1 oscillatory expression, we used a statistical method developed by Phillips et al. (Phillips et al., 2017). Timeseries of bioluminescent or fluorescent HES1 were first detrended to account for long-term trends. We used a squared covariance detrending approach with lengthscale 7.5h, consistent with previous guidelines (Phillips et al., 2017). The detrended timeseries were then analysed using Gaussian Processes (GP). Briefly, the GP approach uses experimental timeseries to assesses whether the signal is better described by a non-oscillatory fluctuating or oscillatory covariance model. The approach contains an internal calibration of white noise detector followed by a stringent 5% false discovery rate statistical test to determine oscillatory timeseries. For the bioluminescence data we used the variance of background timeseries to measure detection noise. For the fluorescence data this is not sufficient, as nuclear signal also contains auto-fluorescence, and here we used a modified approach which infers the variance of noise from the fluorescent data by optimising the relationship between the LLR and signal-to-noise ratio as previously described (Manning et al., 2019). GP fitting producing null parameter estimates were excluded. As an additional quality control, we only report as oscillatory timeseries with a minimum of 2 peaks per track. These additional exclusions affected less than 15% of data per experiment.

### Fluorescence Correlation Spectroscopy

Primary *HES1*^*mScarlet-I/mScarlet-I*^ NSCs were plated in 35mm glass bottom dishes (Greiner) (22K-32K cells per square centimetre) and transfected with pRRL-UbC-mVENUS:HES1 plasmid. FCS experiments were performed at day 1 or day 2 post transfection using a Zeiss LSM880 Inverted Airyscan with a Fluar 40x/1.30 Oil M27 objective while the cells were kept at 37oC and 5% CO2. The fluorescent signals were collected on the same track where mVENUS fluorescence was excited with a 488nm laser and emission was collected between 508-552nm, and mSCARLET-I fluorescence was excited with 561nm laser and emission was collected between 606-677nm. Data from mVENUS/mSCARLET-I double positive cells or mSCARLET-I single positive cells were collected using 3×8sec or 1×15sec runs with 0.05-0.1% 488nm laser power and 0.05% 561nm laser power, which were optimised to reduce bleaching. Traces with large spikes/drops in the count rate and bleaching >20% were excluded. Results were comparable between the long acquisition sequences and triplicate short sequences. FCS auto-correlations were then analysed individually with the PyCorrFit software (Muller et al., 2014). Autocorrelation curves were fitted to one-component diffusion model with triplet state and the Levenberg-Marquardt minimisation algorithm was applied with a number of neighbouring data points j=3. The structural parameter (SP) was fixed at 4. The autocorrelation curves with the best model fitting were used and a counts per particle cut-off of 0.05KHz and 0.1KHz for the mVENUS and mSCARLET-I particle estimation respectively was applied.

### Statistical analysis

Statistical analysis was performed in GraphPad Prism 8.4.1. Data was tested for normality with a Shapiro-Wilk test to inform the choice of a parametric or non-parametric statistical test. The lower and upper hinges of the boxplots correspond to the first and third quartiles. No outliers were removed. The statistical tests, sample sizes, number of biological replicates and p-values are reported in each figure legend.

