## Supplemental Information for "HES1 protein oscillations are necessary for neural stem cells to exit from quiescence"

**Supplementary Information**

**Supplementary Figure 1: *Id* gene expression and HES1 dynamic expression profile in proliferative, quiescent and reactivated conditions.**

A) SOX2 and PAX6 immunofluorescence staining of E13.5 LUC2:HES1 LGE NSCs. B) qPCR analysis for *Id1*, *Id2*, *Id3* and *Id4* mRNA expression in E13.5 LUC2:HES1 NSCs in proliferative, quiescent and reactivated conditions. Fold change expression is relative to *Id1* mRNA levels in proliferative conditions (error bars represent standard deviation, n=3 biological experiments). C) Box plots representing the mean square displacement (MSD) for each cell tracked in proliferative, quiescent and reactivated conditions. This is defined as the square of the distance of the cell from the starting point of tracking plotted against the relative time since the beginning of tracking (dots represent full-length individual cell traces, black horizontal lines represent median, number of cell traces analysed: 67 proliferative from n=6, 61 quiescent from n=4, 120 reactivated from n=5 biological experiments). D) Average length of tracking for each cell in proliferative, quiescent and reactivated conditions (dots represent full-length individual cell traces, black horizontal lines represent median, number of cell traces analysed: 90 proliferative from n=8, 61 quiescent from n=4, 120 reactivated from n=5 biological experiments).

**Supplementary Figure 2: HES1 level does not change as NSCs transition from proliferation into quiescence**

A) Graph showing the relative fold change of median luminescence expression between proliferative and quiescent conditions per experiment. Luminescence expression was estimated by the median luminescence expression per cell trace and per condition (error bars represent standard deviation, two-tailed paired t-test, ns=not significant).

**Supplementary Figure 3: DNA sequence of the *Hes1* locus in the *HES1^mScarlet-I/mScarlet-I^* transgenic mice**

A) A DNA sequence encoding for a linker protein, a 3xFlag epitope and the mSACRLET-I protein has been inserted downstream of the last *Hes1* exon and before the 3’UTR.

**Supplementary figure 4: Ectopic sustained HES1 expression does not increase total HES1 level above physiological range**

A) Correlation of HES1:mSCARLET-I or UbC-mVENUS:HES1 fluorescence intensity with HES1:mSCARLET-I or UbC-mVENUS:HES1 concentration per cell from 3 replicate experiments. E13.5 HES1:mSCARLET-I NSCs were transfected with UbC-mVENUS:HES1. HES1:mSCARLET-I fluorescence intensity and HES1:mSCARLET-I concentration was estimated from untransfected cells while UbC-mVENUS:HES1 fluorescence intensity and UbC-mVENUS:HES1 concentration was estimated from transfected cells in the same culture (Pearson correlation, the shaded gray area represents 95% confidence interval) B-C) Box plots showing total HES1 concentration in untransfected (left panels) and UbC-mVENUS:HES1 transfected (right panels) E13.5 HES1:mSCARLET-I NSCs. In the reporter transfected cells, the total HES1 concentration is the sum of UbC-mVENUS:HES1 concentration (ectopic HES1) and HES1:mSCARLET-I concentration (endogenous HES1) depicted on separate box plots. In B) all concentrations have been estimated experimentally by FCS (apart from the HES1:mSCARLET-I concentration in transfected cells which was inferred). In C) all concentrations have been inferred (black horizontal lines represent median, number of cell analysed in B) untransfected= 75, transfected=38, n=3 biological replicates, in C) untransfected=316, transfected=77, n=3 biological experiments)

**Supplementary Table 1**

List of primers used for qPCR related to Methods section.
