## Supplemental Figures for "HES1 protein oscillations are necessary for neural stem cells to exit from quiescence"

### Slide 1
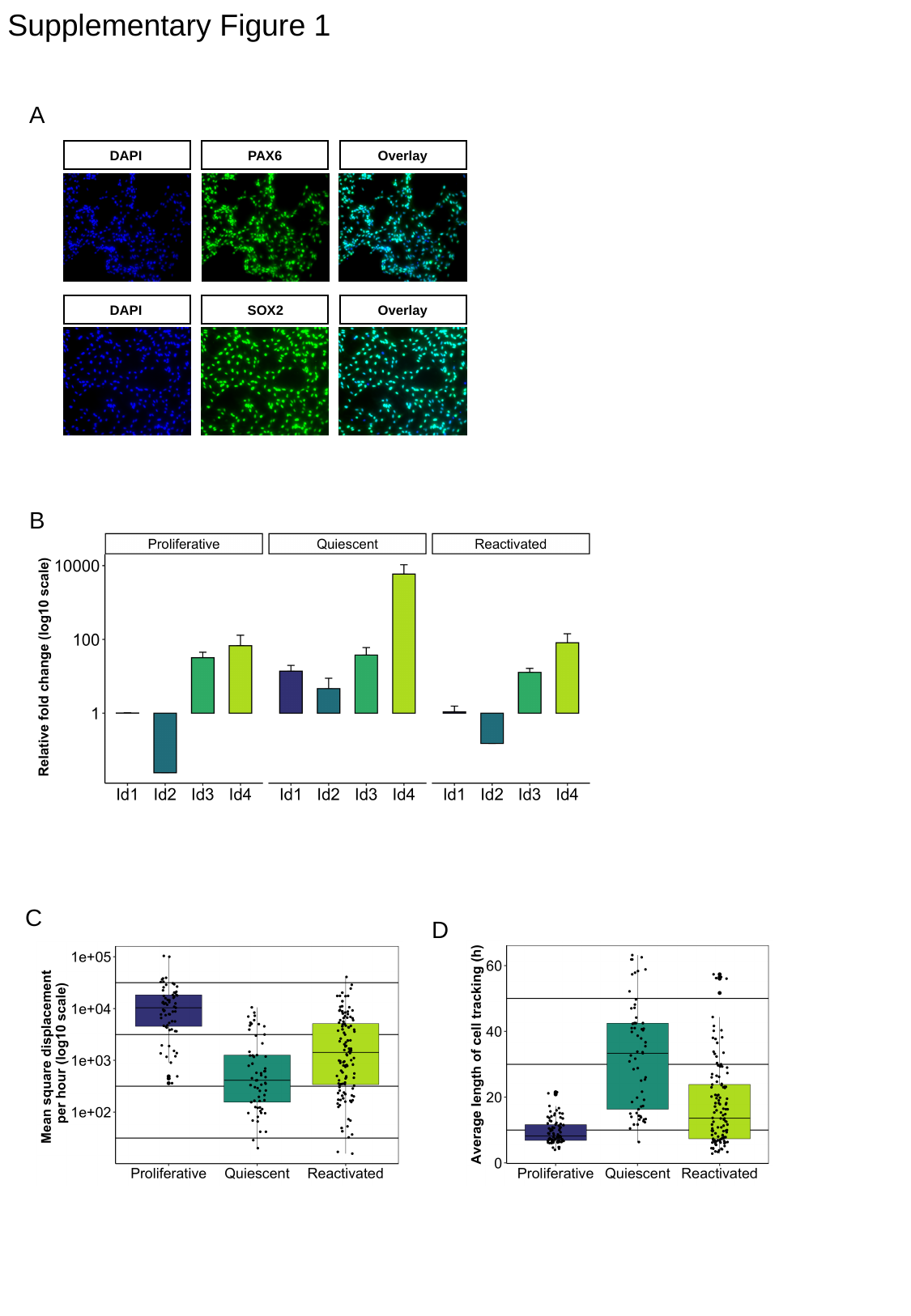

Supplementary Figure 1
A
DAPI
PAX6
Overlay
DAPI
SOX2
Overlay
B
C
D

### Slide 2
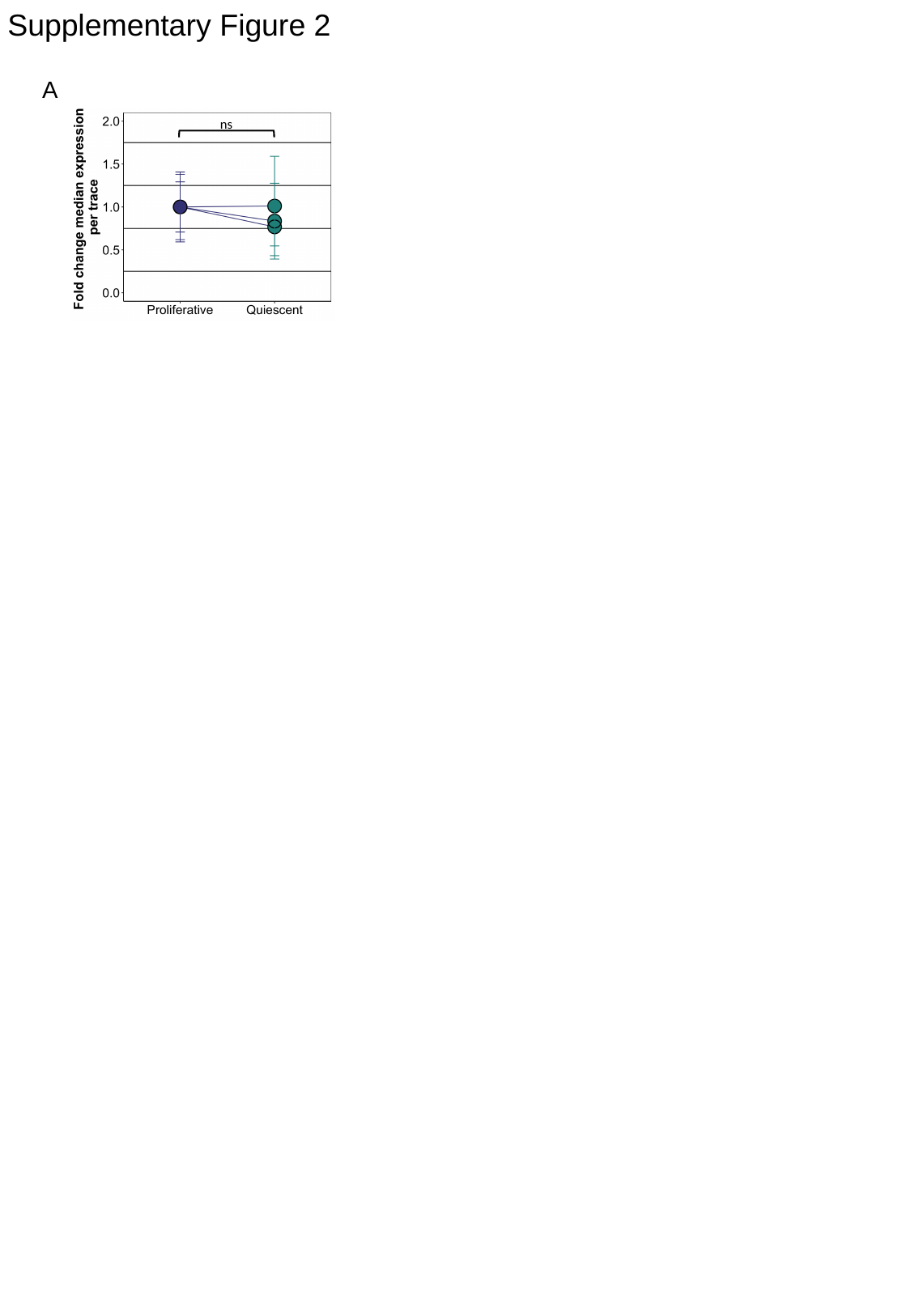

Supplementary Figure 2
A
ns

### Slide 3
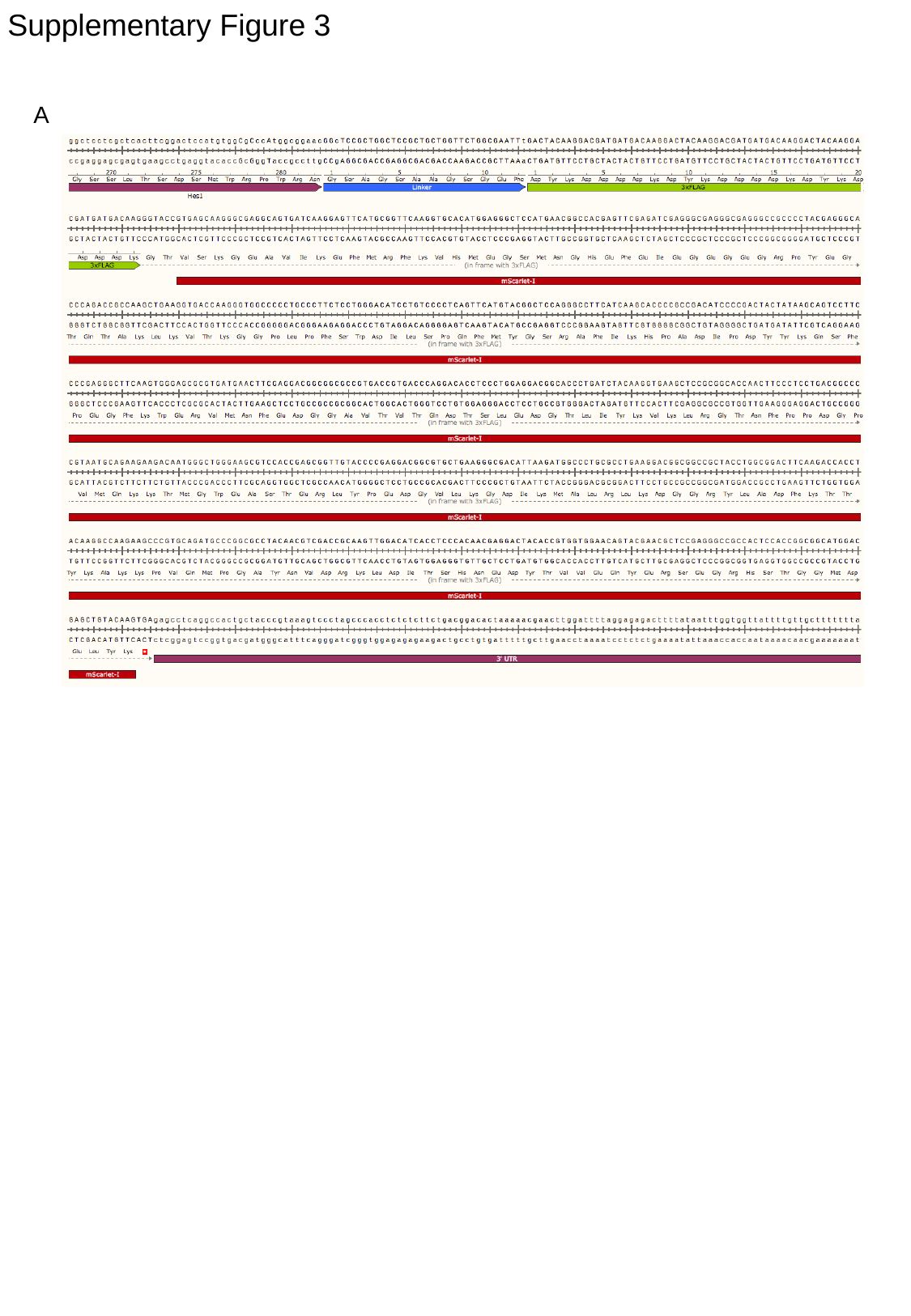

Supplementary Figure 3
A

### Slide 4
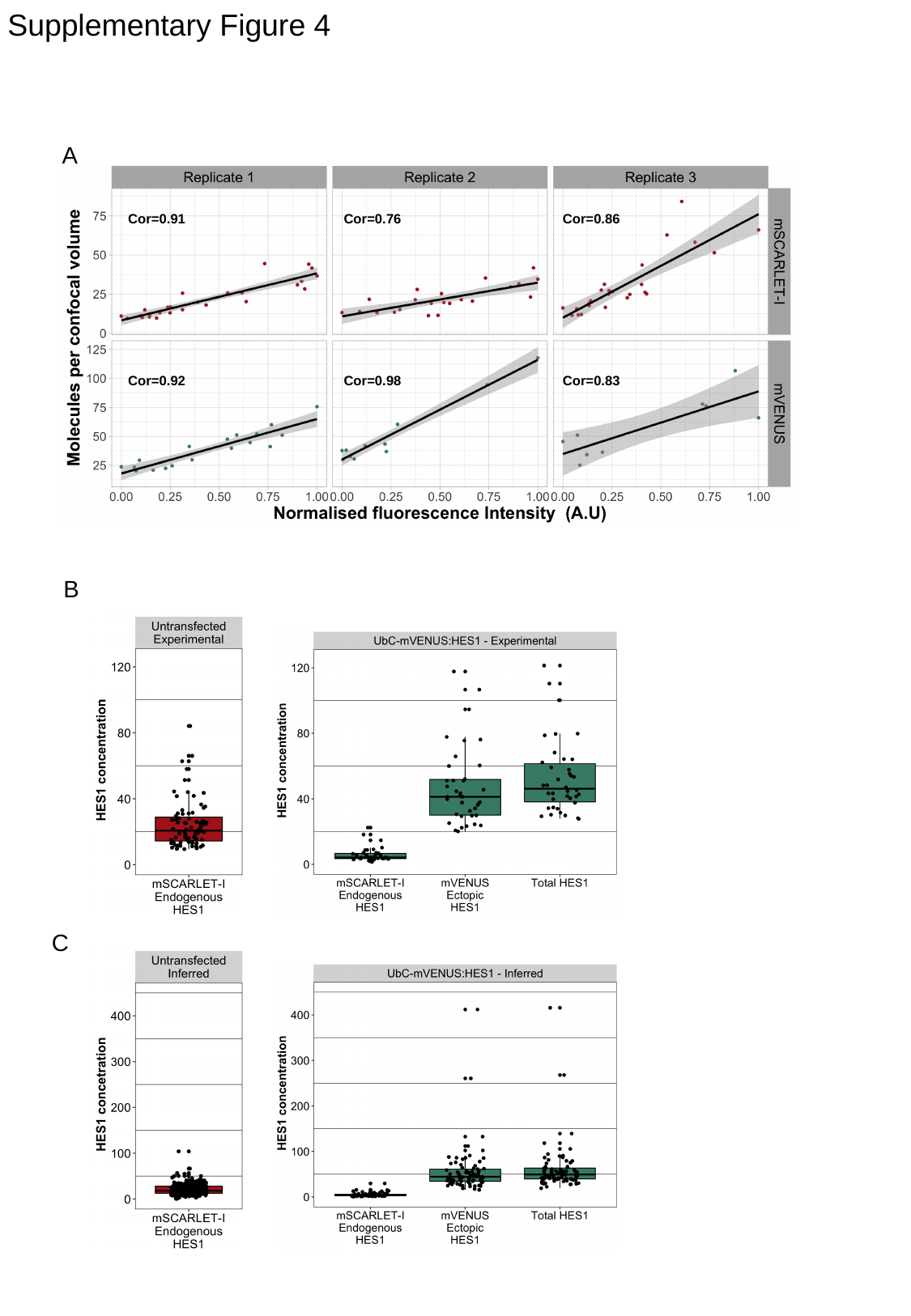

Supplementary Figure 4
A
Cor=0.91
Cor=0.76
Cor=0.86
Cor=0.92
Cor=0.98
Cor=0.83
B
C

### Slide 5
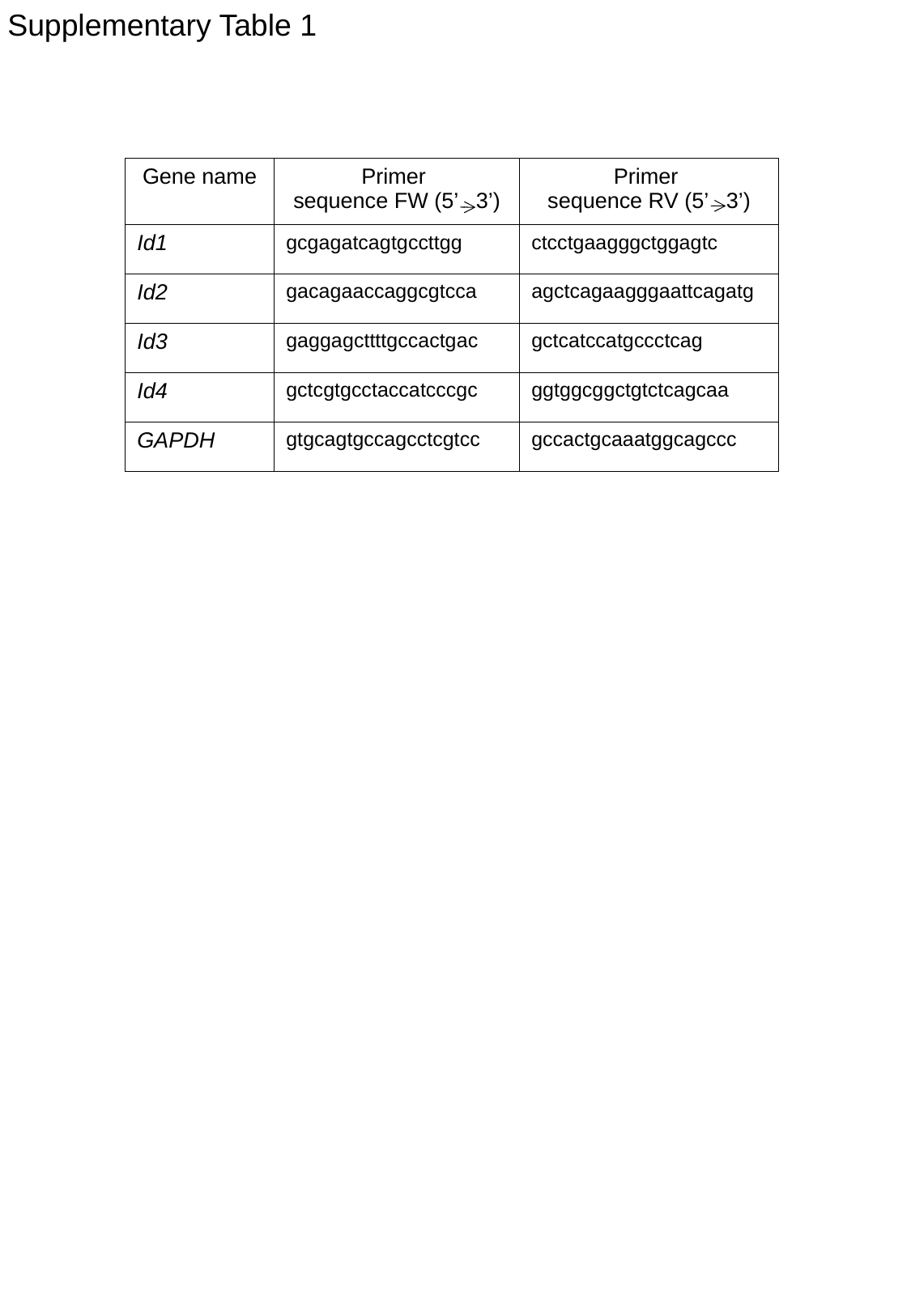

Supplementary Table 1
| Gene name | Primer sequence FW (5’ 3’) | Primer sequence RV (5’ 3’) |
| --- | --- | --- |
| Id1 | gcgagatcagtgccttgg | ctcctgaagggctggagtc |
| Id2 | gacagaaccaggcgtcca | agctcagaagggaattcagatg |
| Id3 | gaggagcttttgccactgac | gctcatccatgccctcag |
| Id4 | gctcgtgcctaccatcccgc | ggtggcggctgtctcagcaa |
| GAPDH | gtgcagtgccagcctcgtcc | gccactgcaaatggcagccc |
